# From affect to control: Functional specialization of the insula in motivation and regulation

**DOI:** 10.1101/102368

**Authors:** Tor D Wager, Lisa Feldman Barrett

**Affiliations:** Columbia University; Boston College

## Abstract

The insula plays a key role in a wide range of brain processes, from viscerosensation and pain to motivation, emotion, and cognitive control. While human neuroimaging studies in all these domains report activations in the insula, little systematic attention is paid to anatomical subdivisions that may provide the basis for functional sub-regions. We conducted a meta-analysis of insular tasks across studies in four domains: emotion, pain, attention switching, and working memory. Using a priori subdivision of the insula based on anatomical studies, we provide evidence that different sub-regions are preferentially activated in different tasks. We suggest that the ventral anterior insula is most important for core affect, a term that describes broadly-tuned motivational states (e.g., excitement) with associated subjective feelings. The dorsal anterior insula, by contrast, may be critical for developing and updating motivational states with specific associated actions (i.e., goals). This region is activated by cognitive control tasks, pain, and some tasks that elicit affective processing. The posterior insula, including SII and portions of parietal operculum, is distinctly activated by pain, providing a double dissociation between pain and tasks that elicit emotions.

---

In 1953, scientists found a new way to think about the human mind. The computer metaphor cast the brain as the ultimate machine, the *deus ex machina* of conscious will reduced to cogs and wheels cranking away behind the curtain, and a new discipline—cognitive science—was born. The computer metaphor continues to be pervasive: information is maintained online in short term memory ‘buffers,’ addressed and tagged, and written to permanent long-term memory storage. Attention works to enhance ‘information processing.’ Reasoning and even some forms of mental imagery proceed through propositional, symbolic representation. In the heyday of the cognitive science revolution, emotion was regarded as a volume knob, an orphaned sideshow in the great circus of the machine.

New developments in recent years are beginning to challenge this view of the mind. The cognitive and neural sciences have increasingly merged, and researchers studying the neural bases of thought and behavior have found a surprising degree of overlap between the brain regions engaged in cognitive and affective processes. The anterior cingulate and insular cortices, among other regions, are engaged reliably in animal and human studies of both central topics in cognitive science—working memory, long-term memory, control of attention—as well as tasks designed to isolate emotional processes. Understanding behaviors or feelings at a given instant in time is a task that requires consideration of the whole organism, and this challenge has forced us to reexamine the ways in which we think about cognitive and emotional processes.

There are two main views on the relationship between cognition and emotion. One view, which has existed in various incarnations since the time of the ancient Greeks, is that cognition and emotion are embodied in two separable, opposing systems. Feeling emotion turns off cognition, and thinking dampens emotional impact (Drevets & Raichle, 1998; Mayberg et al., 1999; Metcalfe & Mischel, 1999; Mischel, Shoday, & Peake, 1989).

Another view seems initially to stand in opposition to the first: Emotion is critical to motivating cognition and behavior, and the roots of attentional control lie in affect. According to this view, emotions arise from cognitive appraisals of situations (R. S. Lazarus, 1991; Richard S. Lazarus, 1991b; Scherer, Schorr, Ed, & Johnstone, 2001; C. A. Smith & Ellsworth, 1985; Craig A. Smith & Lazarus, 2001), which particularly involve evaluations of how objects and events affect the self. The basic affective experience that arises when a self-relevant event occurs has been labeled “core affect” (J. A. Russell & Barrett, 1999). Core affect is the seed of full-blown emotion, and from affective responses arise the physiological and motivated response tendencies that have been shaped over the course of our evolution to promote adaptive cognitions and behaviors. Thus, in this view, emotion and cognition are not opponents in a zero-sum tug of war. Rather, they are synergistic partners in the game of adaptive self-regulation, each shaping the direction of the other.

The answer may be that neither of the metaphors of opposition and synergy is adequate. Emotion does not stop cognition; rather, it directs cognition into channels most appropriate for the situation. In threatening situations, attention is focused on the perceived threat, and extraneous thought stops. In safe situations dominated by positive affect, the impulse to explore and build new skills may prime a broad repertoire of thoughts and behaviors (Fredrickson, 2001). We are at the frontier of exploring the physical brain systems that give rise to thoughts and feelings, and many of these questions may be addressed empirically.

Understanding the roles of certain key brain structures may provide critical information on how affective information shapes attention, and how thoughts direct, or in some cases stem, the flow of affective signals in the brain. Some of the most important such regions are likely to be those that lie at the physical junctions between neocortical and evolutionarily older subcortical nuclei—the limbic and paralimbic regions, so named because they form a limbus, or border, around the oldest parts of the brain (Maclean, 1955, 1958; Papez, 1995). Cortical limbic and paralimbic areas are generally thought to include the cingulate cortex, parahippocampal gyrus and entorhinal cortex, orbitofrontal cortex, and the insula.

## The insula: A key link between cognition and affect

In this paper, we focus on the insula as a potential nexus for motivated cognition and emotional behavior. The insula has long been considered part of the emotional and viscerosensory brain (Janig & Habler, 2002; Maclean, 1955), with multiple roles in regulating physiological and psychological homeostasis (Flynn, Benson, & Ardila, 1999). The insula and surrounding operculum contain primary cortical representation of smell and taste (Francis et al., 1999; Rolls, 1996), viscerosensation (Craig, 2002), and pain perception (Coghill, Sang, Maisog, & Iadarola, 1999; Davis, Kwan, Crawley, & Mikulis, 1998). For this reason, it has been termed “limbic sensory cortex” and associated with the subjective feeling of emotional states, or the “feeling self” (Craig, 2002, 2003). Recent evidence from neuroimaging studies corroborate this view. The insula is commonly activated in emotion tasks, predominantly those associated with negative or withdrawal-related emotions (Phan, Wager, Taylor, & Liberzon, 2002).

Several individual examples illustrate that the insula plays a broad role in the development of subjective, self-relevant feelings. In a recent study, Sanfey and colleagues (Sanfey, Rilling, Aronson, Nystrom, & Cohen, 2003) found insular activity when participants received an unfair monetary offer from what they believed was another human—but not when the offer came from a computer. Why does the same loss sting more if it results from another’s intention? It could be because the loss also signals social rejection, “unfairness,” and/or the motivating possibility of reprisal or control of the situation. Another study found that pain-responsive portions of the anterior insula showed decreased activity when a placebo—a medication with no real effect, but believed by participants to be a potent analgesic—was administered prior to pain (Wager, Rilling et al., 2004). A third study showed common activations in the insula when feeling pain and when observing a loved one experience pain (Singer et al., 2004).

However, the story suggested by these studies, that of the insula as a structure for *feeling*, is incomplete. It omits a host of studies that suggest alternative roles for the insula—in particular, a role in cognitive control. If the anterior insula is the seat of emotional awareness, why is the anterior insula (as we show here) activated in so many cognitive tasks ostensibly devoid of emotion (Wager & Smith, 2003)?

One possibility is that cognitive tasks simply activate a separate portion of the insula and frontal operculum devoted to attentional control. Without detailed comparison of results from cognitive and affective tasks, we cannot tell. Another possibility is that these cognitive tasks, which share as a common feature the requirement for executive control of attention, share common psychological processes with affective tasks. These could include a role for motivated decision making in goal formation, updating of task relevance based on affective information, and affective error-detection processes in cognitive tasks.

A second unresolved issue is the relationship between pain and affect, and how that relationship is informed by insular participation in both. Does pain involve affective representations, as is commonly believed (Melzack & Casey, 1968; Rainville, Duncan, Price, Carrier, & Bushnell, 1997), and do the insular regions involved in “pain affect” form the basis for the “feeling self”? Again, we can ask the same questions about whether pain and emotion activate common parts of the insula, or whether common involvement of the “anterior insula” in pain and emotion is in fact an overgeneralization.

These questions are approached using meta-analysis of human neuroimaging studies (Fox, Parsons, & Lancaster, 1998) across four domains: emotion, pain, attention shifting, and working memory. We begin with functional subdivisions suggested by cytoarchitectural and functional studies in animals (Figure 1), and ask whether these subregions produce meaningful dissociations between tasks in human neuroimaging studies.

**Figure 1.**
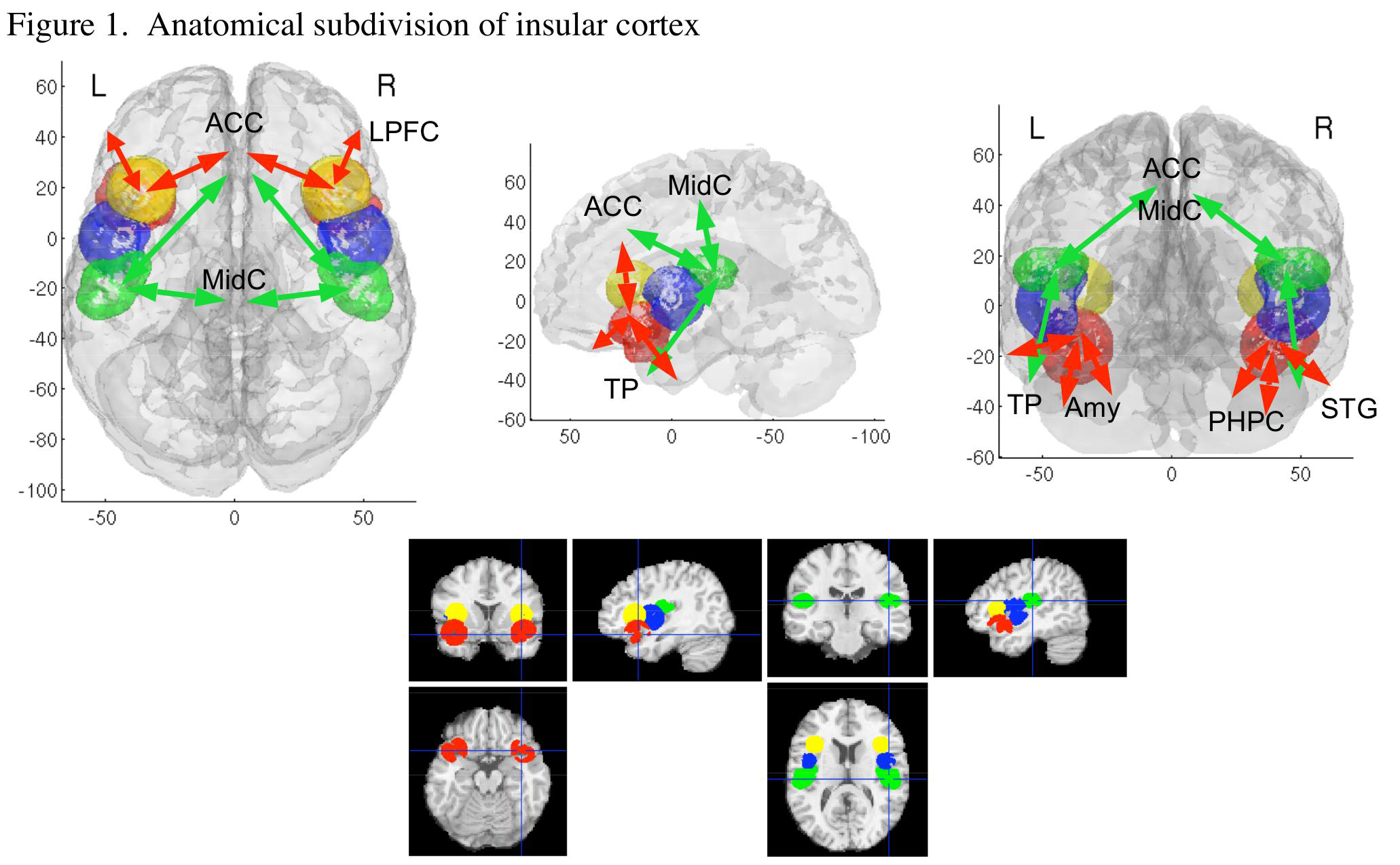
Anterior ventral agranular (Ag) is shown in red. Dorsal anterior dysgranular and adjacent frontal operculum (Adg) are in yellow. Mid-insula (Mdg) is in blue. SII and adjacent parietal operculum are in green. Arrows show a schematic of projection patterns for posterior and anterior regions. ACC, anterior cingulate; MidC, mid-cingulate; LPFC, lateral prefrontal cortex; TP, temporal pole; Amy, amygdala; PHPC, parahippocampal cortex; STG, superior temporal gyrus

We then ask whether activation of particular insular subregions, or the pattern of activation across subregions, provides useful information about which psychological task or process elicited the activations. This is a fundamentally different question that is typically investigated in neuroimaging studies to date. Rather than asking, “What brain areas implement attention shifting,” we ask, “Is there a pattern of activations that uniquely identifies attention shifting, and separates it from other processes?”

We present evidence for distinct insular sub-regions, and we use an inductive approach to develop hypotheses about the processes they may implement. In this paper, we argue for a distinction between pain and emotional feelings, which activate the dorsal and ventral portions of the anterior insula, respectively. The pattern of activations we observe also suggests that attentional control tasks activate a specific part (dorsal anterior) of the insula, in common with pain.

We frame our conclusions in terms of hypotheses to be tested. One hypothesis is that dorsal anterior insula is directly involved in attentional control, and pain activates this region because it recruits mechanisms of executive attention (Eccleston & Crombez, 1999). Alternatively, we present the hypothesis that executive attention recruits anterior insula because this region links general motivational tendencies with specific action plans—i.e., it is involved in goal formation and re-formation. The ventral anterior insula may represent motivational states with very general action tendencies (e.g., affiliate, protect), and the dorsal aspect represents motivational states associated with specific action plans. This distinction is consistent with patterns of results across all the task domains we studied.

## Methods

### Functional subdivisions of insula

Figure 1 shows a digitized parcellation of human insular cortex and the overlying operculum into four distinct regions based on the cytoarchitectural and fiber tract tracing studies of Mesulam and Mufson (Mesulam & Mufson, 1982a, 1982b; Mufson & Mesulam, 1982). Regions were defined electronically on the single-subject Montreal Neurological Institute (MNI) template (Evans et al., 1992) and masked to include gray matter or voxels within 3 mm of gray matter using custom software in Matlab 6.5 (Mathworks, Natick, MA). Each region is discussed in turn below.

**Agranular insula / prepyriform cortex (red).** The agranular insula— Ag, shown in red, and so named for its indistinct laminar structure and apparent lack of stellate (granule) cells prominent in cortical input layers—is the part of insula that surrounds the prepyriform cortex, the primary site of olfactory input to the cortex. It spans medial portions of the temporal pole, caudal orbitofrontal gyrus, and ventral anterior insula.

This portion of the insula and adjacent cortex responds to primary odor reinforcers in humans and animals (Critchley & Rolls, 1996; Dade, Zatorre, & Jones-Gotman, 2002) and appears to play a role in the representation of drive states (e.g., hunger, Freeman). It is most closely connected to the medial orbitofrontal cortex, a primary function of which may be updating the reward value associated with stimulus cues (Baxter, Parker, Lindner, Izquierdo, & Murray, 2000; Rolls, 2004; Wallis, Dias, Robbins, & Roberts, 2001), and also sends and receives projections from the mid-insula, the pericallosal anterior cingulate, and the medial temporal lobes (red arrows in Figure 1). According to Craig (2002), it is via the connection to OFC that the anterior insula has its effects on the valenced property of core affect.

Evolutionarily, Ag and immediately adjacent portions of temporal pole and orbitofrontal cortex were developed for advanced chemoreception, particularly the association of behavioral states (approach/eating and avoidance of noxious environments or foods) with particular chemosensory representations. Thus it earned it the title of prepyriform cortex or primary olfactory cortex, although its selectivity for odors and/or tastes remains under debate. Its evolutionary origins suggest that agranular insula may be part of a core of a system for evaluating primary reinforcers and determining appropriate motivational states—that is, core affect. It is expected to be particularly involved when internal states (like hunger) drive valuation, or when stimuli in the environment signal a direct link to positive (reward), negative (threat) value, or other motives such as self-protection or affiliation (Fridja). Based on this analysis, we expect this subregion to respond during tasks that induce feeling states in people—that is, tasks that elicit emotions.

**Anterior dysgranular insula (Adg, yellow).** The superior portion of the anterior insula (yellow in Figure 1) is dysgranular, with incomplete laminar structure and a cytoarchitectural appearance intermediate between agranular paleocortex and fully developed neocortex (Mesulam & Mufson, 1982a). It blends into the fully-laminated frontal operculum. Although it has not been well differentiated from other parts of anterior insula, this region is commonly activated in tasks that require executive control of attention, including those that require manipulation of information in working memory (Wager & Smith, 2003), response inhibition (Nee, Jonides, & Wager, 2004), and shifting attention (Wager, Reading, & Jonides, 2004). However, its role in these tasks has been underappreciated, perhaps due to the emphasis in the literature on affective and autonomic facet of anterior insular function (Critchley, Wiens, Rotshtein, Ohman, & Dolan, 2004; Phillips et al., 1997). Its proximity to agranular insula and interposition between this older structure and the lateral prefrontal cortex—thought by many authors to be the seat of executive control over attention and action (Miller, 2000)—suggest that it may be important for translating undifferentiated drive states into specific action plans.

**Mid-insula (blue).** The middle portion of the insula (blue in Figure 1) is also dysgranular, and it is connected primarily with neighboring areas of insula.

**SII and parietal operculum (green).** The superior bank of the posterior insula contains SII, primary sensory cortex for pain and itch (Craig, 2002, 2003; Craig, Chen, Bandy, & Reiman, 2000). Its major bidirectional connections are with parts of the ventromedial nucleus of the thalamus, which transmit sensory input from the body (Craig, 2002); multiple regions of the anterior and mid-cingulate (Mesulam & Mufson, 1982b; Mufson & Mesulam, 1982), consistent with known cingulate motor regions in the monkey (Picard & Strick, 1996); adjacent S1 and sensorimotor cortex; and parts of the anterior temporal cortex.

### Study selection and meta-analysis

Studies for meta-analysis were compiled using Medline and ISI Web of Science database searches, and from additional references found in papers, and are drawn from the literature on pain, emotion, working memory, and attention shifting. Although we do not claim that the studies here represent an exhaustive list, we have tried to be as inclusive as possible. We analyze activations (not deactivations) by tabulating peak coordinate locations reported in studies of healthy participants (excluding patient studies), in keeping with more extensive reports in our previous meta-analyses (Phan et al., 2002; Wager, Phan, Liberzon, & Taylor, 2003; Wager, Reading et al., 2004; Wager & Smith, 2003).

Many studies reported multiple independent comparisons, or contrasts—for example, an emotion study might compare perception of fear faces vs. a neutral baseline, and perception of angry faces against a separate set of neutral images. In practice, many studies compared activations in multiple conditions to the same baseline; this strategy, and the fact that the same sample of participants is used across comparisons, prevents these contrasts from being truly independent, but we treat them here as if they were. The relevant data used for meta-analysis was whether each independent contrast within each study activated each anatomical region of interest.

Analyses included chi-square analyses, k-nearest neighbor (KNN) classification, and rule-based classification using activation in single areas or pairs of areas to predict tasks. Chi-square analyses test whether contrast counts within a region differ among task conditions, controlling for the overall number of contrasts studied in each condition. This analysis is similar to those reported in our previous studies, with one improvement: by counting contrasts, rather than peaks or studies, we can better account for studies with multiple contrasts while avoiding overweighting of studies that report many peaks. More methodological detail can be found in our other papers (Phan et al., 2002; Wager, Reading et al., 2004; Wager & Smith, 2003).

KNN analysis was performed on the presence/absence of regional activations in each anatomical subregion across different types of tasks. In practice, we formed an indicator matrix, the rows of which were contrasts, and the columns of which coded for both tasks and regional activations. This matrix is the building block of the multiple correspondence analysis framework, which can be used for multivariate analysis of categorical data (Bouilland &. Loslever, 1998). Task conditions (e.g., approach-related emotions and withdrawal-related emotions) were coded in one set of columns, and the presence of activation in each region was coded in another set of columns. Ones indicated that the contrast belonged to a task condition or activated a region, and zeros indicated lack of membership or activation.

As task categories in the present analysis were mutually exclusive, the task indicators were recoded into a class vector that described the task performed in each contrast. KNN analysis analyzes the *k* nearest neighbors (we chose a relatively standard value of 3) to each point in the data space, and uses those to form a prediction about the task type of each point. In our analyses, each contrast was assigned a task classification based on the known task classifications of the three contrasts that produced most similar activation profiles across the eight insular subregions. Errors in classification were calculated by comparing the known classes with estimated classes (summarized in a confusion matrix). We chose to derive several summary measures of classification accuracy: the hit rate, or correct classification rate for each class; the false alarm rate, or rate at which other task types are classified as a particular task; and a sensitivity measure (A’) based on the combination of hit rate and false alarm rate (Swets, 1988). When only two classes are compared, a single A’ gives the discriminability of the two classes. For more than two classes, each class has its own A’, reflecting the discriminability of that class from the others.

A particular concern with classifiers is that it is relatively easy to construct a classifier that will perfectly predict classes in a particular dataset, but cannot generalize to new data because it uses information that is too specific to the particular dataset. A KNN classifier with k = 1 is an example, as each task will always be given the class neighbor of it’s one nearest neighbor – itself. Larger k helps to avoid this problem. An additional procedure is cross-validation, which divides the set into a larger training set and a smaller test set. The training set (85% of the data in our analyses) is used to derive classifications, and classification of a smaller test set (15%) is performed using similarities between each test contrast and the training data. We divided the dataset into training and testing sets multiple times (200 in the present analyses) and estimated test classes, hit/false alarm rates and discriminability for each sample, providing a cross-validated estimate of the true ability of the KNN algorithm to correctly classify contrasts into task categories according to their insular activation patterns.

Rule-based classification was performed in a rudimentary fashion in this paper by simply considering the probability that a study belonged to a particular task category, given activation in each subregion individually and a limited range of task alternatives. This can be written as *p*(task=T | activity), where *p* denotes posterior probability given the observed data, *task* is used generally to refer to psychological condition or class, and *activity* signifies the presence of an activation peak in a particular region of interest.

In a Bayesian framework, the posterior probability estimate *p*(task=T | activity) = *p*(activity | task=T) * *p*(task=T) / p(activity). The first term, the likelihood of activity given task T, is the proportion of contrasts of task type T that activate the brain region. The second term, the prior probability of a task being of type T, could be calculated for a particular sample based on the frequency of tasks in the database; but we wanted to generalize to a world in which all tasks could be studied equally frequently, so we assign p(task=T) = 1. p(activity) is the sum of percentages of contrasts activating across task types, and normalizes the posterior probability estimates across tasks to 1. This normalization excludes the possibility that a task belongs to none of the set of possible tasks in the analysis. Thus, the task with the maximum posterior probability estimate is simply the best choice among relative alternatives.

We report posterior probabilities in the Results as percentage scores that reflect predictions of task condition given a) observed activity in single regions or combinations of two regions, and b) the set of alternative tasks included in the analysis. In comparing pain with emotion tasks, for example, observing bilateral SII activity may be associated with a 100% posterior probability for pain, but that probability estimate will change if additional task types beyond pain and emotion are considered. We leave the full development of probability estimates into full classifiers, considering all regions, for future work.

## Results

### Pain and feelings elicited by emotional stimuli activate distinct subregions of insula

SII activation was strongly associated with painful stimulation (Figure 2), but not with emotion. Activation in either right or left SII was associated with a 90% posterior probability of pain, and bilateral SII activation with 100% probability of pain (no emotion studies activated bilateral SII). Mid-insula activation in either hemisphere was associated with an 80% probability of pain stimulation.

**Figure 2.**
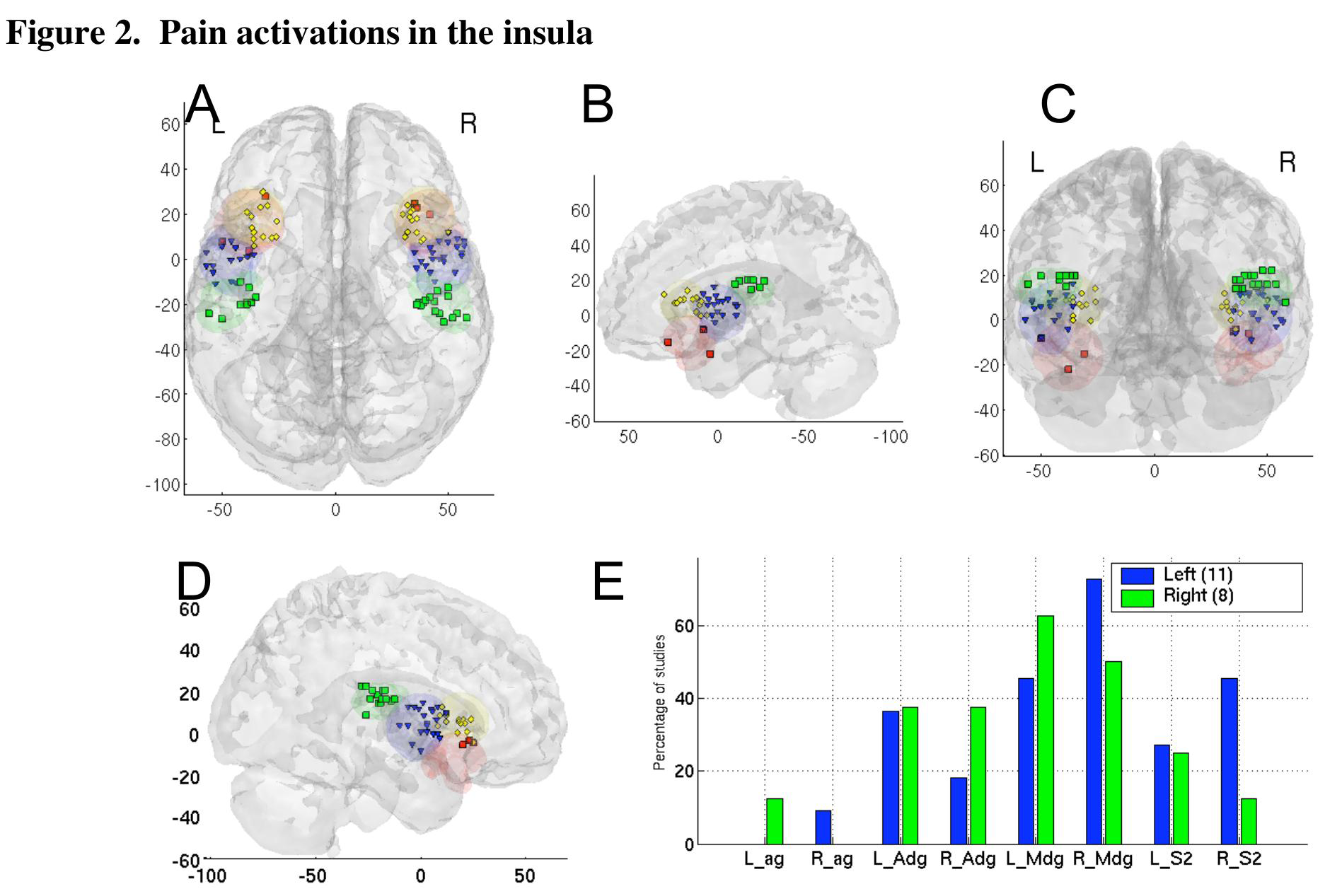
A-D) Points on the transparent brains reflect pain activation coordinates in insular subregions: green for SII, blue for mid-insula, red for anterior agranular (ventral) insula, and yellow for anterior dysgranular insula (superior). E) Activation counts by independent contrasts for painful stimulation by region (x-axis) and side of body stimulated. Bar heights reflect the proportion of contrasts within each body side that activated each region.

Agranular insula was likewise relatively strongly associated with emotional processing. Left agranular insula was associated with a 77% probability of doing an emotion task, with 60% for the right agranular subregion. The only pain studies to activate bilateral agranular insula were Bense et al. (Bense, Stephan, Yousry, Brandt, & Dieterich, 2001), the only study to use vestibular pain, and (Becerra, Breiter, Wise, Gonzalez, & Borsook, 2001). Right agranular insula was reported by Brooks et al. (Brooks, Nurmikko, Bimson, Singh, & Roberts, 2002), using thermal pain on the left arm. There was also a bias toward left agranular insular activation in emotions (Figure 3E) that did not hold for frontal opercular and dysgranular region, in contrast with Craig’s (Craig, 2003) view of the right anterior insula as mediating the ‘feeling self.’

**Figure 3.**
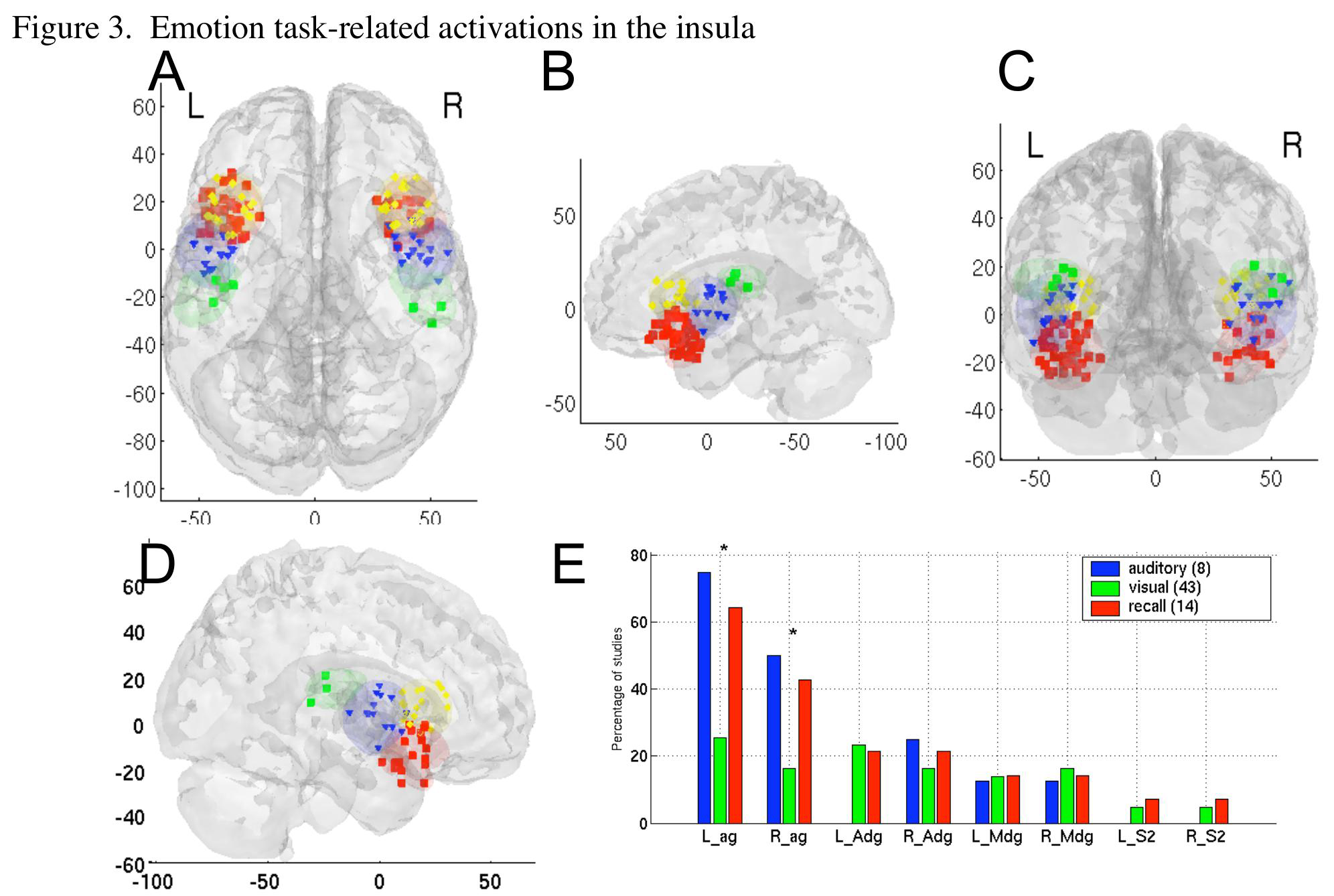
A – D) Glass brains showing peak activation coordinates for emotion tasks. E) Contrast counts by type of material eliciting emotion, as in Figure 2. * indicates significant differences across conditions for a region

Classification results indicate that pain tasks and those that involve emotion are relatively discriminable based on their patterns of activation. KNN classification resulted in a cross-validated correct classification of 91% for emotion tasks and 52% for pain tasks, with a high discriminability (A’) of 2.17.

However, to view emotion tasks as ‘of a piece,’ or as a natural kind, is an extremely limited view. Rather, we would like to move towards defining the processes within particular emotion tasks that produce activations in the agranular insula. Indeed, additional analyses showed that emotional activations were unevenly distributed across emotion induction methods (Figure 3E). Both auditory induction (voices, screams, etc.) and recall-induced emotions produced substantially more frequent activations in agranular insula than did visual inductions.

Comparison of approach and withdrawal related emotions—broadly construed, happiness and anger vs. sadness, fear, and disgust—provided weak support for a dorsal/ventral distinction. Although chi-square results for individual areas were not significant, approach was associated with agranular activation in each hemisphere, whereas withdrawal was associated with superior agranular activation in each hemisphere. Analysis by valence, which included anger as a negative emotion, produced yet less consistent results.

Activation in individual regions was not highly predictive of approach vs. withdrawal state (the highest was right anterior dysgranular insula, associated with a 73% probability of withdrawal), but activation of both left and right anterior dysgranular or both left and right mid-insula led to 100% withdrawal classification. Thus, no studies of approach activated either of these subregions bilaterally. However, the probability that a withdrawal study produces these activations is also low (6%, or three studies (Damasio et al., 2000; Phillips et al., 1997; Simpson et al., 2000)). Damasio et al. studied emotion-induced recall of personal events, and found activations in all portions of the insula except SII. Phillips et al. studied perception of disgust faces during a gender identification task. Simpson et al. showed this activation in response to aversive pictures during concurrent number judgments.

Was superior insula activated by the aversive quality of the emotions, or by cognitive demand associated with performing concurrent cognitive tasks? Virtually all studies activating this region required cognitive demand (chi2 = 5.24, p <.05 in left Adg and 2.04, n.s. in right Adg), with the exception of Canli1998, who showed activation in right Adg. Right Ag showed significantly more frequent activations with no cognitive demand (chi2 = 6.11, p <.05), indicating that the few studies of visual and auditory passive perception that activated Ag tended to do so in the right hemisphere. Recall-induced emotions involve cognitive demand de facto, as well as eliciting emotion, but they did not frequently activate Adg. Thus, cognitive demand seems to be a more powerful predictor of Adg activation than does the aversive quality of the emotion elicited. Analysis by individual emotion produced no notable distinctions in activation of insular subregions.

This pattern suggests that the agranular insula may play a role in the experiential component of emotions—the process of inducing and feeling an emotion, and perhaps experiencing associated motivational tendencies, rather than simply perceiving emotional stimuli. An alternative is that subvocal verbalization or language plays a primary role in aurally and recall-induced emotions, given the role of insula in understanding and producing language (Habib et al., 1995). However, as we discuss below, analysis of verbalizability of materials in working memory tasks revealed no effects on agranular insula.

The pattern of results contrasts sharply with that found in the amygdala, which in previous meta-analyses showed selectivity for visual stimuli, particularly perception of fearful faces, suggesting a role for the amygdala in the visual perception of threat (Murphy, Nimmo-Smith, & Lawrence, 2003; Phan et al., 2002). Interestingly, individual studies have reported deactivations in the amygdala and agranular insula during pain (Derbyshire et al., 1997).

### Shifting attention and executive working memory activate anterior dysgranular insula

Attention shifting results, summarized in Figure 4, show a clear cluster of peaks only in the left and right superior anterior insula. Most of these peaks lie at the junction of the dysgranular insula and frontal operculum. Although percentages are not high overall (25% of switching studies activated this insular subregion), the consistency of their location suggests that the result is reliable. Additionally, several studies showing activation in agranular and mid-insula produced peaks right at the border of the anterior dysgranular region, suggesting that the borders of this area do not quite capture the observed pattern. Analyses by type of switching (i.e., among locations, tasks, objects, rules, or attributes of objects) yielded no consistent results.

**Figure 4.**
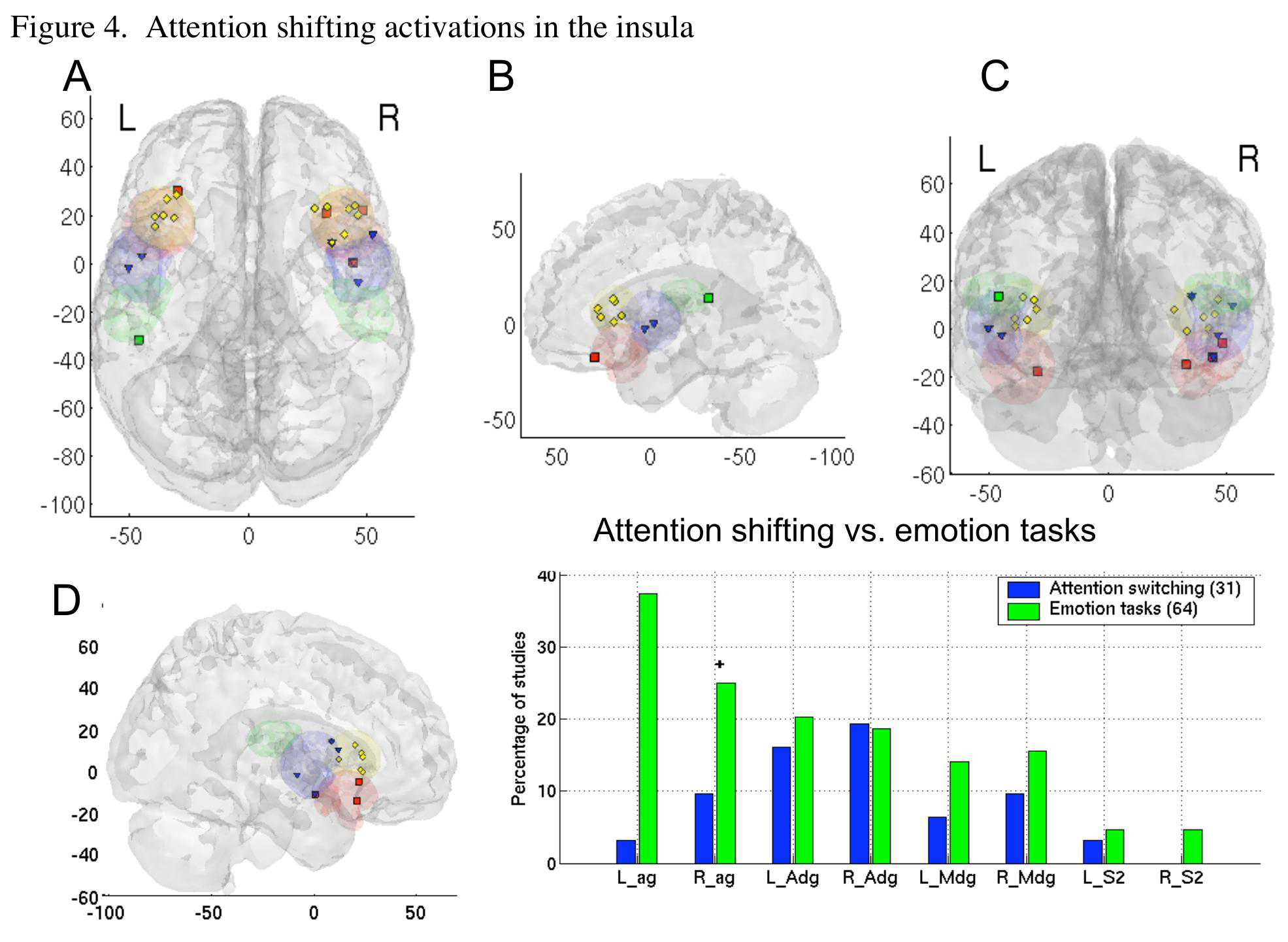
A-D) As in previous figures. E) Counts for attention compared with emotion tasks. Axes are as in previous figures. * indicates significant differences across conditions for a region at p <.05.

Comparing switching results to results from emotional tasks (Figure 4E), we observed that there was significantly more agranular insula activity in emotional contrasts. Although emotion tasks and shifting tasks activated anterior dysgranular insula about equally frequently, KNN classification was able to accurately distinguish switching from emotion tasks, with cross-validated correct classification rates of 60% for switching and 78% for emotion, and a discriminability A’ of 1.41.

These results demonstrate that executive attention activates a subset of insular regions activated in emotion tasks. One possible conclusion is that elicitations of emotion involve re-directions of attention, and thus attentional control is a component process engaged when emotions are aroused.

A similar profile of activations was found for executive working memory (Figure 5), with one notable exception. Working memory tasks produced consistent activation in right agranular insula, overlapping with emotional task activations. Activation of both this region and the right anterior dysgranular region were significantly more frequent for tasks involving executive control of working memory, suggesting that right agranular insula is affected by executive demand. Because the activation profile was otherwise similar as that for switching attention, KNN classification did not discriminate well between switching and working memory tasks, with correct classification rates of 23% for switching and 81% for working memory, and a low A’ of –1.41, which suggests worse-than-chance performance.

**Figure 5.**
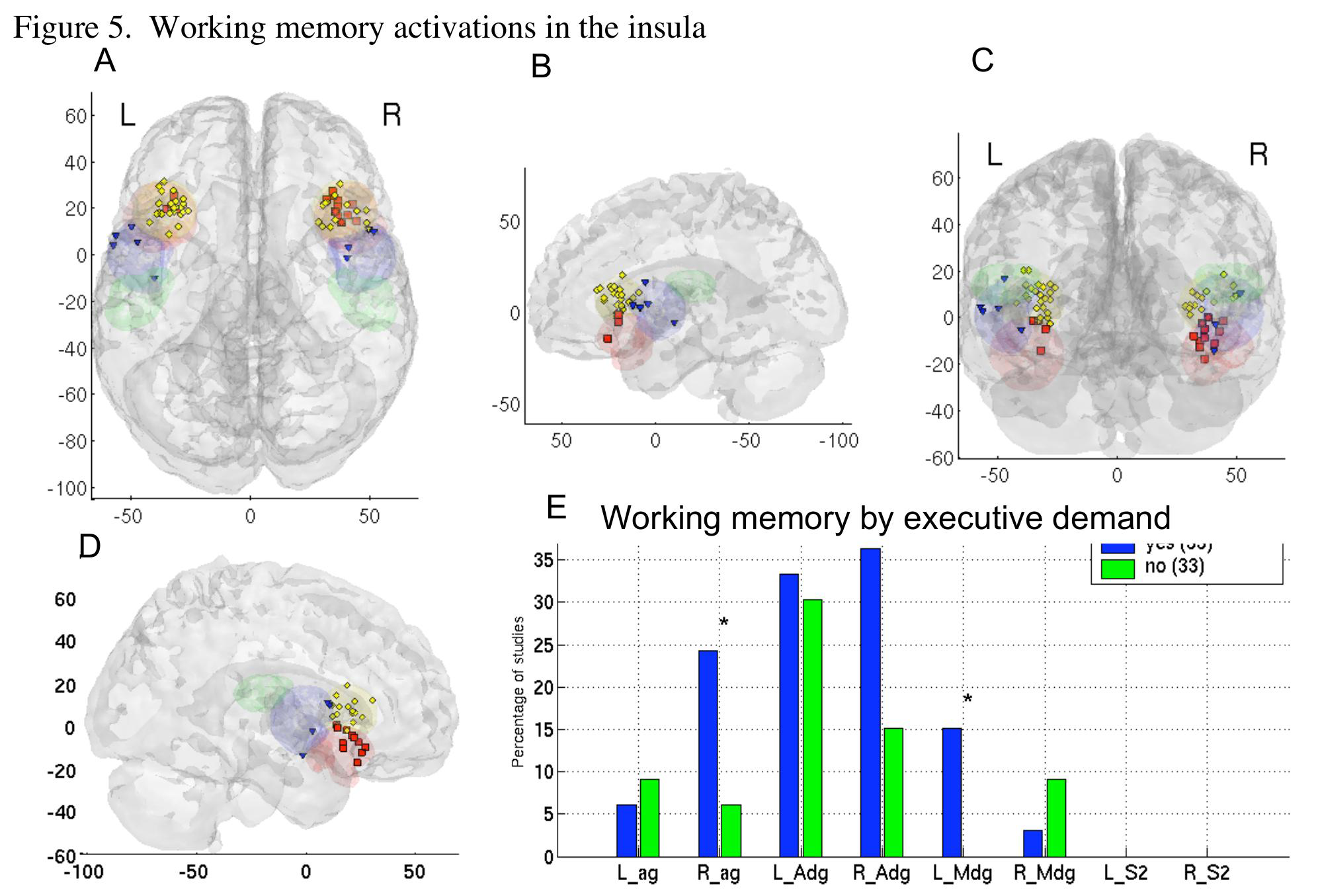
A-D) As in previous figures. E) Counts for by executive demand on working memory. Axes

## Discussion

Our results suggest that the ventral anterior agranular insula is activated consistently by neuroimaging studies involved in aurally and recall-generated emotion induction, particularly in the left hemisphere. Executive manipulation of information in working memory also engages the right agranular insula, but pain and attention shifting do not. We suggest that a key function of the agranular insula may be in representing afferent homeostatic information from the body, for the purposes of subjective evaluation. This process is central for generating sets of motivated responses—and, as we discuss below, is closely tied to the psychological concept of core affect (James A. Russell, 2003). These findings contradict older broad conceptualizations of the emotional brain, which suggest that the right hemisphere is the more emotional, or that the left hemisphere is more selective for positive emotions and the right for negative ones (although this latter may apply to affective styles and specifically to the lateral frontal cortex, as reviewed in (Wager et al., 2003)).

Our findings also suggest that the anterior insula can be subdivided into two parts: the ventral, agranular part discussed above, and a superior dysgranular part that is contiguous with the frontal operculum (Mesulam & Mufson, 1982a). The localization of attention switching, working memory, and pain results specifically to the superior anterior insula supports this distinction, as does the mass of activations from emotional recall tasks in the ventral anterior insula. If anterior insula represents the ‘feeling self’ (Craig, 2002), we may ask which part of the anterior insula is most critical. Our data suggest the agranular portion is most critical, particularly on the left side—in contrast to Craig’s argument that the interoceptive self is localized to the right anterior insula (Craig, 2002). Furthermore, pain produces only superior, agranular activation, suggesting that the interoceptive process for pain is different than for emotion; pain affect is not the same as affect per se.

One possible explanation of the ventral-dorsal distinction is that emotional recall tasks (ventral) elicit a very broad sense of emotionality, with broad motivational/action repertoires, whereas pain (dorsal) carries affective signals that motivate very specific escape or avoidance action repertoires. Demand on executive control of attention (dorsal) may also motivate action-specific changes in behavior: When a task is difficult, error signals are generated that the organism is doing the wrong task or doing the right task in a substandard way. Attention must be reallocated or the strategy changed. This is at the heart of “mental effort,” and it is accompanied in cognitive tasks by autonomic reactions. What is “substandard” must be determined by evaluating the anticipated benefits and harms of current performance with respect to the self. Thus, the ventral anterior insula is central to broad feelings such as “happy” or “sad” that lead to general strategies, and dorsal anterior insula is central to specific affective signals that lead to specific strategies—“this action is wrong; change it.” Both are essentially processes of valuation, requiring assessment of benefit and harm to the self, and they lead to core affective/motivational states that proscribe response patterns. We discuss the concepts of both valuation and core affect below.

Finally, the parietal operculum (SII) is critical for somatic signals that impact homeostasis—it is activated relatively uniquely by sensory pain (and related somatic sensations). SII is particularly involved in processing pain, and is perhaps involved more directly in representing somatic and visceral autonomic input (Craig, 2002, 2003). However, almost no studies of emotion or attention control activate this subregion. One of the oldest theories of emotion is that, to relative degrees, we develop emotion by interpreting autonomic signals ascending from the body. An extreme form of this theory is the James-Lange theory—that we “are scared because we run (or sweat, or our heart races).” If SII represents somatic interoceptive signals, then our results suggest that somatic interoception—at least at the relatively early stage of SII processing—is not important for emotion. However, it must be noted that interoception itself has produced activations in various parts of the insula, particularly the anterior portion (Critchley et al., 2004), suggesting the need for more precise localization of viscerosensory functions across task contexts (Cameron, 2001; Cameron & Minoshima, 2002; Critchley, Melmed, Featherstone, Mathias, & Dolan, 2001; Critchley et al., 2004).

The mid-insula was activated by some studies in all domains, but was not activated by a high percentage of studies in any domain except pain. Our results do not show any clear, convincing role for the mid-insula that is not more characteristic of another subregion, so we restrict the bulk of our interpretations to the more diagnostic subregions.

In the remainder of the discussion, we elaborate on two key psychological constructs, valuation and core affect, that seem to be related to anterior insula function in particular.

**Valuation.** Organisms continually judge situations and objects for their relevance and value – that is, whether or not their properties signify something important to well-being. A number of studies strongly suggest that such evaluations occur automatically, continuously, and often subconsciously (J. A. Bargh, 1990; J. A. Bargh, Chaiken, Govender, & Pratto, 1992; J. A. Bargh, Chaiken, S., Raymond, P., & Hymes, C., 1996; Chaiken, 1993). Objects and situations rarely have intrinsic value or meaning; rather, value is acquired through cognitive appraisal of their significance and assessment of their impact on well-being (Clore, 2000; R. S. Lazarus, 1991a, 1991b). Thus, valuation is a process that depends on comparison of the external and internal worlds, and valuation is central to what we mean when we say something involves affect. Affective processing includes those neural processes by which an organism judges, represents, and responds to the value of objects in the world (Cardinal, 2002).

**Core affect.** The products of valuation are motivational states—tendencies to act in particular ways, often linked to the accomplishment of goals. Evolution has shaped valuation processes to produce particular states in particular situations, some highly automatic or ‘canalized’, others more flexible. Learning also shapes valuation, linking particular situations with particular motivational states. These *states* we term core affect (James A. Russell, 2003; J. A. Russell & Barrett, 1999).

In our view, the valuation process continuously updates our core affect. In a sense, everything that has been said about “emotion” may be true of core affect. The hardwiring to support it is present at birth (Bridges, 1932; Emde, 1976; Spitz, 1965; Sroufe, 1979). It can be acquired and modified by associative learning (Cardinal, 2002). It can exist and influence behavior without being labeled or interpreted, and can therefore function unconsciously, although extreme changes that capture attention or deliberate introspection may allow core affect to be represented verbally.

Interpreting our meta-analytic findings in the broader context of human and animal literature, it appears that different subregions of the anterior insula may play different roles in the evaluative/core affective process. One possibility, as we discussed above, is that Ag is critical for subjective feelings, and Adg is important for signaling a need for a specific change in strategy.

A second, related possibility arises from theory on emotion that posits general and specific action tendencies that are core parts of emotion (Frijda, 1988; R. S. Lazarus, 1991; Richard S. Lazarus, 1991b). Frijda (Frijda, 1988; R. S. Lazarus, 1991; Richard S. Lazarus, 1991b), for example, argues that emotion is a process of translating meaning (i.e., valuation) to action readiness, which can be as general as the desire to affiliate that is associated with joy or the desire to engage the world that is associated with excitement, or as specific as the desire to harm a specific person one is angry at with a specific implement that is available at hand.

Thus, Ag may represents what might be termed broad, nonspecific action tendencies associated with general emotions, and Adg may translate affective signals into specific action plans, given the stimuli and choices available in the current situation. Fredrickson (Fredrickson, 2001) has argued that positive emotions are associated with broader action tendencies and negative emotions with narrower ones, and our meta-analytic findings in the Ag and Adg are consistent with this pattern. Both are different stages in the translation of core affect into motivated behavior.

**From motivation to attention.** The machinery allowing one to pay attention is widely thought to involve dorsolateral prefrontal and superior parietal cortices (Cutrell & Marrocco, 2002; Sylvester et al., 2003). We do not dispute this view; but what then is the role of the insula, and core affect, in attention?

With regard to the control of attention, stimuli in the environment (e.g., new task instructions) or internal states (boredom, hunger) can provide feedback that your current state of attention deployment is no longer optimal given your current needs. Changes in how objects and behaviors are valued can be driven by a number of factors, including verbal information from others, changes in associated reward, or changes in the internal state. Whatever the cause, decreases in the value of the current task or state of attention engender shifts in the motivational state, with consequent shifts in the focus of attention. If valuation of stimuli drives attention, we would expect neurons in dorsal attention-implementing systems to show sensitivity to reward value during attention and working memory tasks. As recent evidence demonstrates with increasing certainty, this is the case (Gehring & Willoughby, 2002; Lauwereyns, Watanabe, Coe, & Hikosaka, 2002; Platt & Glimcher, 1999; Shidara & Richmond, 2002).

### Future Directions

What we have tried to do in this paper is blend confirmatory and inductive approaches to understanding structure-function relationships in the brain. Our analyses were confirmatory in that we used a pre-defined set of anatomical subregions and asked if these boundaries, derived from animal studies, produced meaningful distinctions between different types of functional neuroimaging activations. However, this analysis does not mean that the subregions we chose are the optimal ones; adaptive parcellation of anatomical space based on functional activations would be one approach that could address this question, at the cost of sacrificing confirmatory inferential power.

Our analyses were inductive in that we began empirically, by examining the pattern of activations across different tasks, and using the data to form hypotheses about how the psychological constructs relate to one another. Thus, rather than beginning with the assumption that pain invokes emotion, we asked whether brain responses to pain and emotion induction share a common neural substrate within the insula (and concluded that they share less than might have been anticipated, single studies notwithstanding (Singer et al., 2004)).

One future direction has just been identified: to derive an anatomical parcellation that, given knowledge about activation of each parcel, gives the best information about the psychological nature of the task being performed. This will require adaptive analysis methods targeted at the right level of anatomical detail: too broad, and the map loses specificity; too narrow, and it is driven by the ideosyncracies of past results and loses the ability to generalize to new studies.

A second direction is to apply new methods of classification to produce accurate mappings between brain activity and psychological function. This must be done, at least using functional neuroimaging, by generalizing across a mass of mappings from psychological function to measured brain activity. This essentially Baysian process must proceed in a meta-analytic framework. We have suggested two types of classifiers here. One is simple rule-based classifiers, of the form [If brain activity in X -> Then Task A]—for example, if bilateral SII activity is observed, then a pain stimulus is almost certainly being perceived. The second is pattern-based classifiers such as KNN that use the pattern of activation and non-activation across all regions. In this study, the first method provided some surprisingly powerful classification rules based on simple presence of activation in a subregion. However, the pattern-based methods are also promising, as they can capture more complex configurations of activation across multiple regions.

More work needs to be done to improve these classifiers and validate them with additional studies. For instance, on the input side, coordinates could be weighted by reliability or quality measures, as well as by the degree to which they are representative of activation in a particular subregion. On the algorithmic side, combinations of rule-based and pattern-based classifiers may prove useful.

The critical point is that the pattern of activations across studies has promise for informing us about the relationships among pain, emotion, perception of emotion, and cognitive control processes—in general terms, about the relationships among psychological constructs. In this sense, the usefulness of neuroimaging data can be tested empirically, by the metrics of consistency, reliability, and predictive power.

